# Morphological comparison between cell-like entities from mammalian tissues and Precambrian microfossils

**DOI:** 10.64898/2026.03.22.713352

**Authors:** Elena Angela Lusi, Federico Caicci, Marco Quartuccio, Claudia Rifici

## Abstract

The fossil record of the Precambrian era preserves some of the earliest evidence of life, yet these ancient microfossils primarily reveal morphology rather than function, leaving unresolved questions about how early cells lived, replicated and evolved. The RNA world hypothesis proposes that primordial organisms relied on RNA for both information storage and catalysis, but direct living systems reflecting such biology remain poorly characterized.

Here we describe cell-like entities isolated from mammalian tissues, measuring approximately 1–3 μm in diameter and exhibiting morphological similarity to a range of Precambrian microfossils. Ultrastructural comparisons reveal a high degree of correspondence with fossil taxa spanning the Paleoproterozoic to Ediacaran intervals (∼1.8 Ga to ∼551 Ma). In addition to these morphological features, the entities display biochemical characteristics, including RNA-dominant nucleic acid content and particle-associated reverse transcriptase activity.

These observations indicate that the cell-like entities described are not inert, but represent biologically active systems. The combined ultrastructural and biochemical features raise the possibility that biologically active entities comparable to those observed in Precambrian microfossils may occur in contemporary biological contexts.

## 1. Introduction

Fossil microorganisms constitute the primary record for understanding the earliest evolution of cellular life and the major transitions in biological complexity. For most of the first four billion years of Earth’s history, the biosphere was dominated by unicellular forms and their fossilized remains provide critical insight into processes such as eukaryogenesis, the emergence of multicellularity, sexual reproduction, cellular differentiation and the molecular innovations that shaped genome evolution (Donoghue, 2020). This record is preserved predominantly within Precambrian sedimentary rocks spanning the Hadean, Archaean and Proterozoic eons (∼4.0–0.54 billion years ago), where microscopic fossils represent the principal evidence for early cellular systems. Many of these microfossils, some exceeding 1.8 billion years in age, have long been interpreted as reflecting biological entities capable of metabolism, replication and ecological interaction.

From the Archaean to the late Proterozoic, the fossil record is characterized by a wide diversity of microscopic forms, including organic-walled vesicles (acritarchs), filamentous and colonial organisms, and complex multicellular or embryo-like structures. Among the most extensively studied are the acritarchs, which appear from at least the Paleoproterozoic and display considerable diversity in wall structure, ornamentation and inferred life cycles. Similarly, the phosphatized microfossils of the Doushantuo Formation have been interpreted as preserving cleavage-stage embryos or embryo-like organisms, exhibiting regular internal partitioning and staged developmental progression. Proterozoic deposits have also yielded fungi-like forms characterized by filamentous growth, cellular differentiation and complex wall architecture. Collectively, these assemblages demonstrate that sophisticated cellular designs—including compartmentalization, morphogenetic patterning and developmental organization—were established well before the Cambrian diversification of macroscopic life (Sankaran, 2016; Javaux et al., 2010; Loron et al., 2019; Javaux and Knoll, 2017; Grey et al., 2014; Javaux et al., 2001; Tang et al., 2019; Xiao et al., 2004; Li et al., 2023; Butterfield, 2018; Harvey, 2023; Moczydłowska, 2011; Knoll et al., 2006; Butterfield, 2000; Schopf, 1993; Tacker et al., 2022; Javaux, 2019; Joyce and Szostak, 2018; Chatterjee and Yadav, 2016; Chatterjee, 2009; Chatterjee, 2016; Chatterjee and Guven, 2014).

The late Precambrian culminates in the Ediacaran Period (635–541 million years ago), immediately preceding the Cambrian explosion, during which animal body plans diversified rapidly. Although the Cambrian fossil record captures an abrupt increase in macroscopic complexity, it was preceded by more than a billion years of microbial innovation, during which the structural and developmental foundations of later life were established.

Despite this morphological richness, the interpretation of Precambrian microfossils remains inherently limited. Fossilization preserves structure far more readily than function, and consequently, the biochemical composition, genomic architecture and physiological behaviour of early cellular systems remain largely inaccessible. As emphasized by Donoghue and colleagues, fossilized cells can illuminate patterns of organization and macroevolutionary context, yet they cannot, in isolation, resolve the molecular nature of the organisms they represent. This limitation leaves a fundamental question unresolved: what were the underlying biological mechanisms of these ancient cells, and to what extent, if any, have their structural or functional programs persisted into the present biosphere?

The RNA world hypothesis provides a conceptual framework for addressing these questions at the molecular level. Originally articulated by Gilbert and further developed by subsequent studies, this model proposes that early life relied on RNA both as genetic material and as a catalytic molecule, preceding the emergence of DNA genomes and protein-based enzymatic systems. Within this framework, early cellular systems would have required compartmentalization, structural boundaries, and replicative enzymes such as RNA-dependent RNA polymerases, as well as reverse transcriptase activity facilitating the transition from RNA to DNA genomes. While the Precambrian fossil record constrains the temporal context of these innovations, it cannot directly reveal the molecular composition of the organisms that expressed them (Chatterjee, 2013; Chatterjee, 2018; Joyce, 2002; Higgs and Lehman, 2015; Saito, 2022; Müller, 2006; Gilbert, 1986).

An unresolved issue, therefore, concerns whether the cellular architectures observed in the Precambrian fossil record represent entirely extinct forms, or whether elements of these designs may have persisted—either as continuous lineages or as conserved developmental programs—into modern biological systems.

In this study, we describe cell-like entities isolated from human and animal specimens that exhibit morphological similarity to a range of Precambrian microfossils, including acritarchs, Doushantuo embryo-like forms and Proterozoic fungi-like structures. These entities, measuring approximately 1–3 μm in diameter, display complex organic wall assemblies, internal partitioning consistent with cleavage-like organization and staged developmental progression culminating in budding-like reproduction. Ultrastructural comparisons reveal a high degree of correspondence with fossil taxa spanning the Paleoproterozoic to Ediacaran intervals (∼1.8 Ga to ∼551 Ma), bridging Mesoproterozoic to Ediacaran evolutionary intervals.

In addition to these morphological features, the cell-like entities exhibit biochemical characteristics, including RNA-dominant nucleic acid content and intrinsic reverse transcriptase activity, consistent with genome architectures predicted by genetic models of early life (Chatterjee, 2013; Chatterjee, 2018; Joyce, 2002; Higgs and Lehman, 2015; Saito, 2022; Müller, 2006; Gilbert, 1986). The convergence of ultrastructural and molecular features suggests that these entities may reflect deeply conserved biological strategies linking early cellular evolution to extant systems.

The identification of such entities in a modern biological context raises important questions regarding the interpretation of Precambrian microfossils, the persistence of ancient cellular architectures and the diversity of present-day unicellular life. By placing these observations within a palaeobiological framework, this study provides a basis for reassessing early cellular evolution and highlights the potential to investigate, in living systems, structural and molecular features previously accessible only through the fossil record.

## 2. Materials and Methods

### 2.1 Biological material and sample processing

Biological material consisted of human and animal specimens collected over an extended period of investigation. All samples were handled in accordance with institutional guidelines and standard laboratory procedures.

Cellular fractions containing the entities of interest were obtained through differential centrifugation followed by sucrose density gradient fractionation. In parallel, sample processing was adapted to avoid conventional filtration steps (e.g., ≤0.2 μm), routinely used in microbiological and virological workflows, which inherently exclude particles in the 1–3 μm size range. Instead, size-permissive separation combined with density-based fractionation was used to retain intact cell-sized structures while reducing background material. This approach enabled consistent recovery of the entities described here and may explain their absence from previous reports based on standard protocols.

This procedure was applied routinely over a period exceeding ten years, resulting in a curated biobank of purified preparations. Independent isolations were performed across multiple experimental series and laboratories, and comparable fractions containing cell-sized entities were consistently recovered.

Purified fractions were either processed immediately or cryopreserved for subsequent analyses. Repeated isolation and recovery from independent specimens, including samples processed at different times, by different operators and in different laboratory settings, demonstrated the stability of the preparations and reproducibility of the procedure.

### 2.2 Electron microscopy

Purified preparations were examined using standard transmission electron microscopy (TEM) techniques. Samples were fixed, processed, and imaged according to established protocols routinely employed in electron microscopy facilities. Ultrastructural analysis focused on overall morphology, wall architecture and internal organization. Multiple independent preparations from different isolations were examined to confirm consistency of observed features. Observations were reproducible across preparations derived from independent specimens. Imaging was performed at the Electron Microscopy Unit, University of Padua.

### 2.3 Nucleic acid extraction and enzymatic treatments

Total nucleic acids were extracted from purified preparations using standard extraction protocols. To determine the nature of the genetic material, aliquots were subjected to enzymatic digestion using DNase and RNase under controlled conditions.

Sensitivity to RNase treatment and resistance to DNase digestion were assessed by agarose gel electrophoresis and downstream amplification assays, allowing discrimination between RNA- and DNA-based genetic components.

### 2.4 Reverse transcriptase activity assays

Reverse transcriptase (RT) activity was evaluated using lysates derived from purified preparations. Reactions were performed under conditions comparable to those used for standard RT assays, with appropriate positive and negative controls.

To assess whether RT activity was associated with the cell-sized entities, matched samples were subjected to filtration through 0.2 μm membranes prior to analysis. Filtered supernatants consistently failed to generate detectable reverse transcription products, whereas unfiltered preparations retained RT activity. This size-dependent distribution indicates that RT activity is associated with the purified entities rather than with filterable retroviruses.

### 2.5 RNA sequencing and sequence analysis

RNA was isolated from purified preparations and subjected to high-throughput sequencing using standard library preparation and sequencing workflows. Independent sequencing replicates were performed to ensure reproducibility.

Initial assembly using conventional genome reconstruction pipelines did not yield contiguous linear genomes. Consequently, analysis was conducted at the level of individual sequence contigs, focusing on length distribution, compositional features, and the presence of conserved functional domains. Reproducible patterns across independent datasets were used to assess the structural organization of the genetic system without imposing assumptions of linear genome architecture.

### 2.6 Reproducibility and data collection

The entities were consistently recovered from multiple independent human and animal specimens, including tumour-derived material, by different operators across distinct laboratories and clinical settings. Across preparations, they were reproducibly isolated, purified, cryopreserved, sequenced, and subjected to functional assays, including re-infection and antigenic characterization. The persistence of these entities across samples, operators, laboratories and experimental conditions, together with the reproducibility of their structural and biochemical features, argues against sporadic contamination or technical artefact and supports their interpretation as stable, coherent biological microorganisms.

## 3. Results

### 3.1 Electron microscopy reveals living unicellular entities with Precambrian-like morphology

Purified preparations obtained by sucrose-gradient fractionation of human and animal specimens consistently yielded fractions containing unicellular agents measuring approximately 1–3 μm in diameter. Transmission and scanning electron microscopy demonstrate that these entities are bounded by a distinctive organic wall assembly (OWA) and exhibit pronounced internal compartmentalization. Notably, the cleavage-like internal partitioning observed is closely comparable to division patterns described in Doushantuo embryo-like fossils and acritarch-type microfossils (Figs. 1–3 and SI). The sequence of developmental stages observed closely parallels those reconstructed from Precambrian fossil assemblages spanning approximately 1.8 billion to 551 million years ago, indicating a strong correspondence between extant and fossilized cellular architectures.

**Figure 1.**
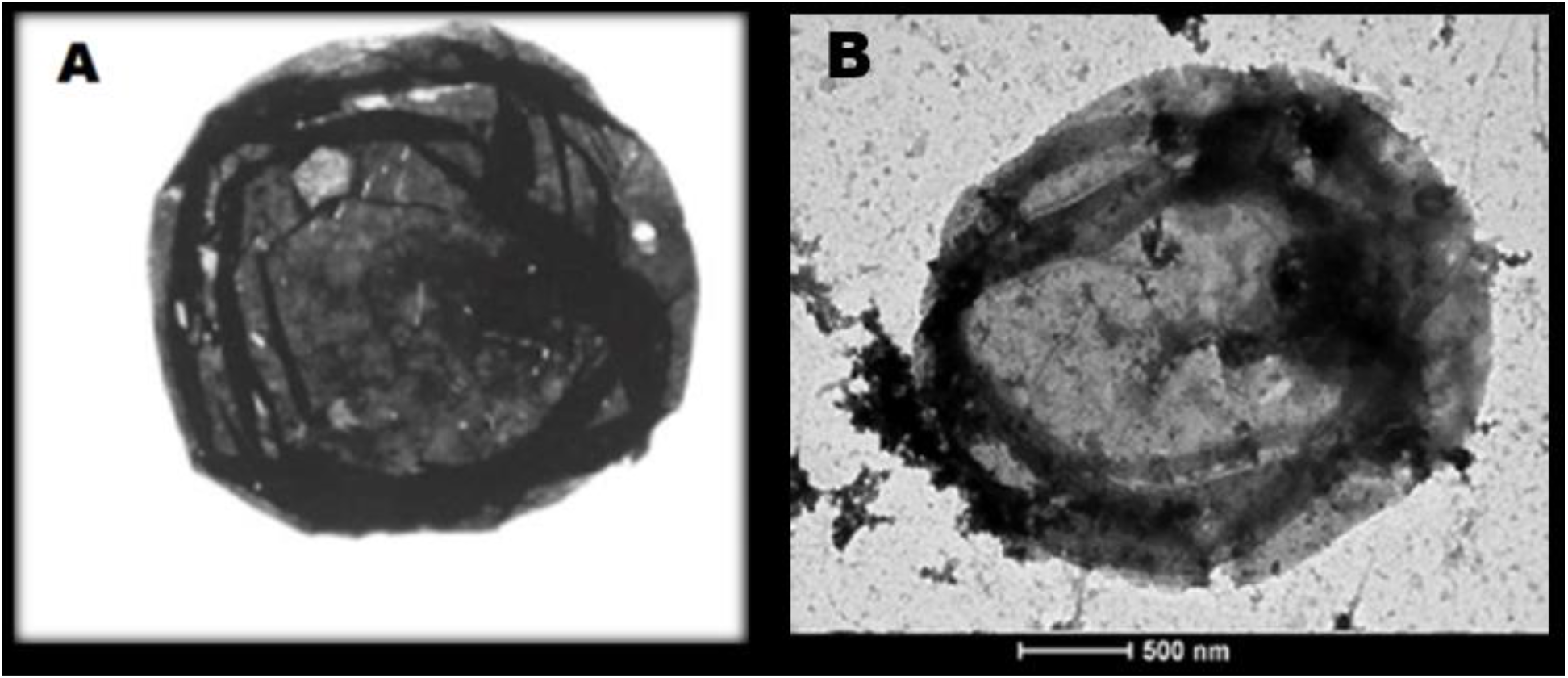
A) Precambrian Acritarchs. B) Unicellular Agent, 3 microns in size, isolated through a sucrose gradient from HPB-ALL leukaemia cells.

**Figure 2.**
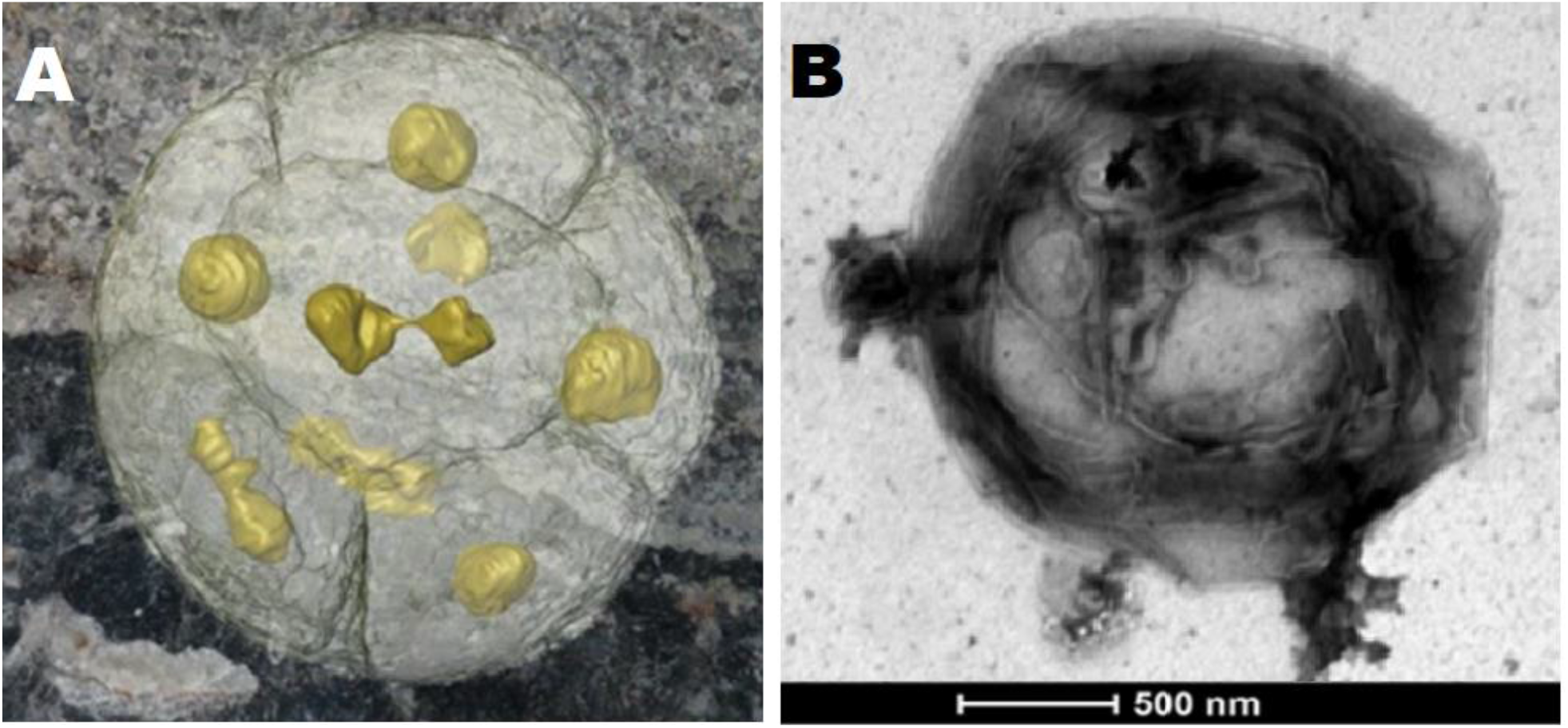
Comparison of Doushantuo fossils with mammalian unicellular entities. **(A)** X-ray microtomographic reconstruction of a fossil from the Doushantuo Formation, China, showing internal partitioning and nucleus-like structures (yellow). (*Enigmatic fossils are neither animals nor bacteria*, Nature 2011). **(B)** Transmission electron micrograph of a living unicellular agent isolated from human cancer tissue. The organism exhibits an organic wall assembly and internal compartmentalization that shows striking correspondence to the partitioned organization observed in the Doushantuo fossil in panel A. Note the similar cleavage-like subdivision and central dense regions.

**Figure 3.**
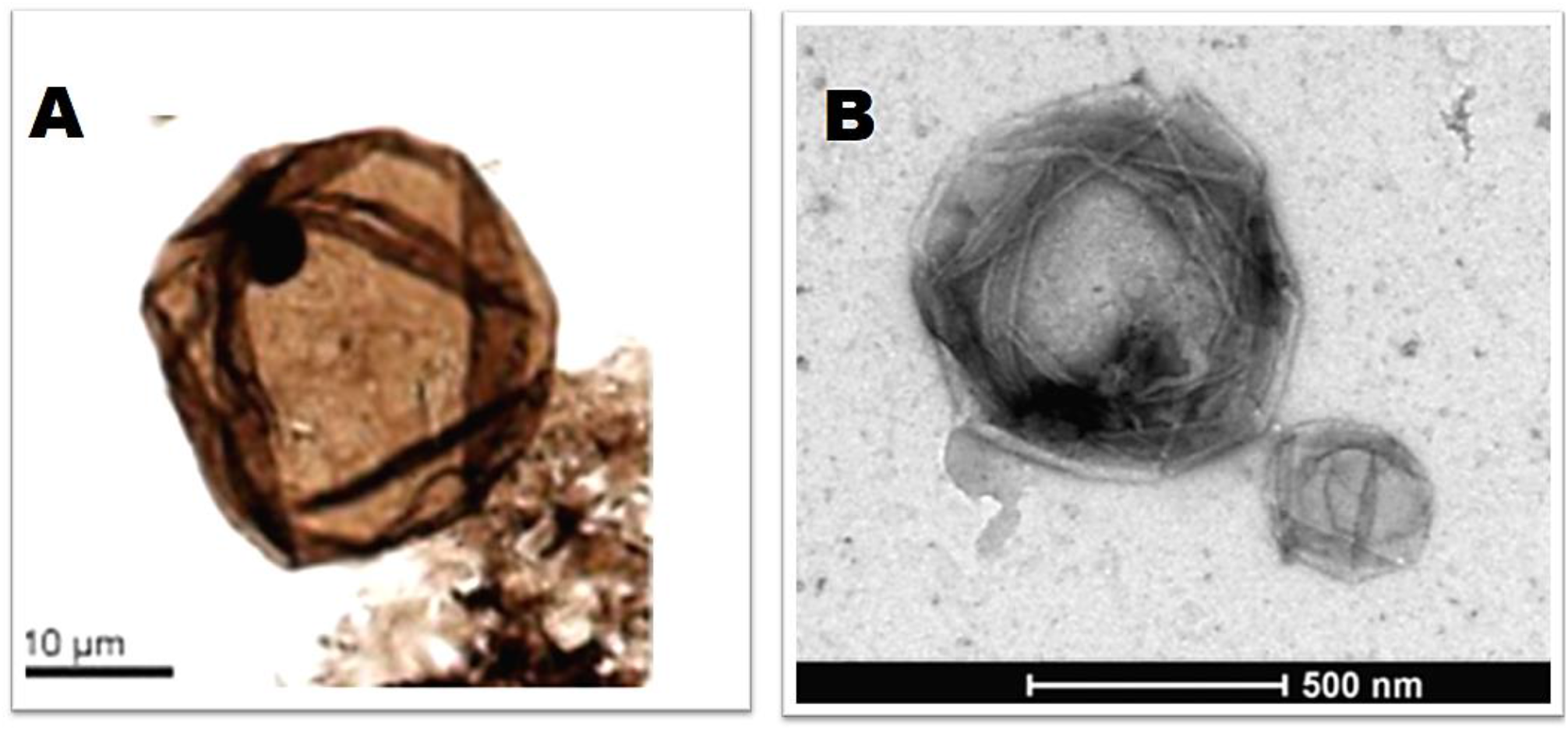
(A) *Leiosphaeridia crassa*, acritarch (incertae sedis) from the lower Shaler Supergroup (Loron et al., 2019), showing a thick-walled vesicle with a central dark inclusion. (B) Living unicellular organism showing a comparable vesicular morphology and internal body (for morphological comparison only).

This correspondence is further supported by comparison with X-ray tomographic reconstructions of Precambrian microfossils reported by Donoghue and colleagues, reinforcing the interpretation that these entities recapitulate fundamental features of early cell-like systems.

On the basis of their morphological congruence with acritarchs and Doushantuo embryo-like fossils, the observed features are consistent with cellular architectures known from the Precambrian eon. The closest fossil analogues span from the Paleoproterozoic (∼1.8 Ga) to the Ediacaran (∼551 Ma). Although direct radiometric dating of living organisms is not possible, the conservation of morphology and developmental patterning supports the possibility that these entities retain deeply ancestral cellular characteristics.

While many observed morphologies closely resemble Ediacaran Doushantuo assemblages, several defining features—including organic wall construction, vesicle-like internal organization, staged maturation and budding- or excystment-like structures—are also characteristic of Mesoproterozoic (>1 Ga) acritarchs. These affinities indicate that the observed architectures are not restricted to late Precambrian forms, but instead reflect cellular designs that likely originated substantially earlier in Earth’s history. Furthermore, in some mammalian tissues, the morphology of these entities overlaps with that of Proterozoic fungi-like microfossils described from Arctic Canada (Loron et al., 2019), which demonstrate that complex, organic-walled unicellular and colonial organisms had already diversified well before the Cambrian explosion, (Fig. 4). Shared features include organic wall assemblies, internal compartmentalization,and staged developmental progression, indicating that similar cellular strategies were established in Precambrian ecosystems. Collectively, these observations support the interpretation that the mammalian-associated entities described here exhibit morphological features consistent with Precambrian microfossils. Their close correspondence to microfossils spanning ∼1.8 billion to ∼550 million years raises the possibility that comparable cell-like systems may remain biologically active in contemporary biological contexts. Representative electron microscopy images are presented in the main manuscript, while a more extensive dataset is provided in the Supplementary Information and deposited in a public repository (Figshare: https://doi.org/10.6084/m9.figshare.31802113).

**Figure 4.**
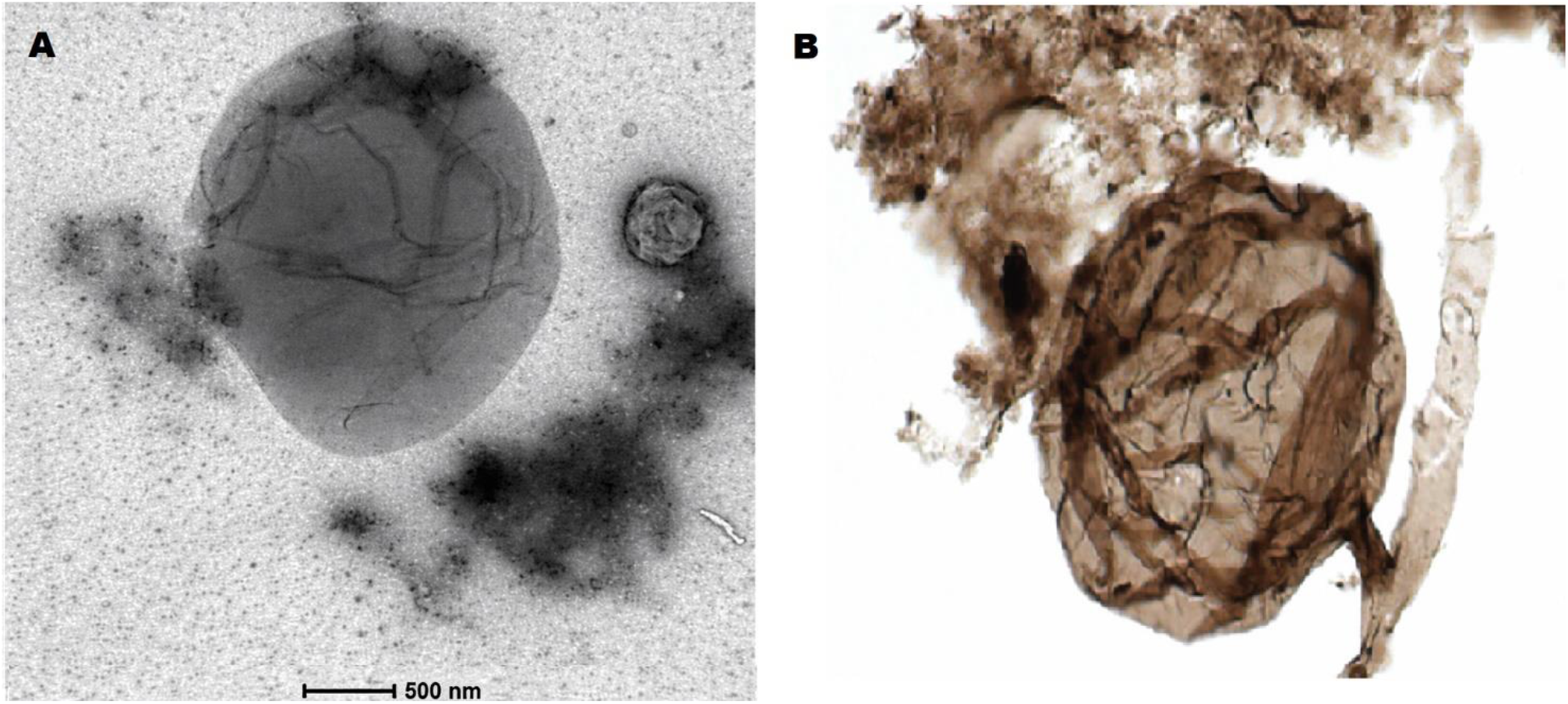
Remarkable morphological similarities between (A) a ∼3 µm living agent isolated from canine transmissible venereal tumour and (B) Precambrian early life microfossils ((Nature. 2019 Jun; 570 (7760):232-235)

**Figure 5.**
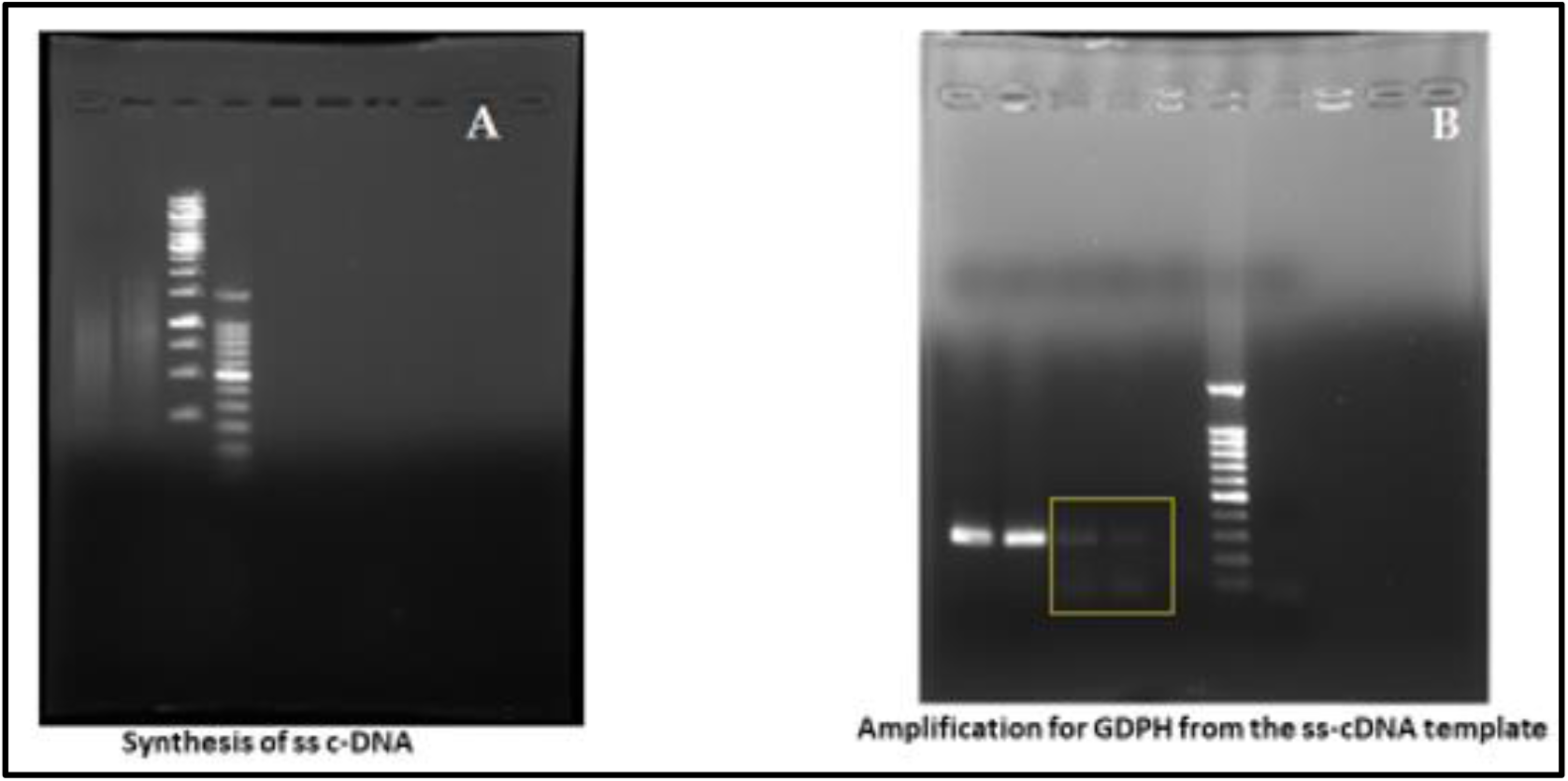
Reverse transcription activity associated with the purified entities. (A) cDNA synthesis detected in lysates derived from the purified preparations (lane 2), compared with a reaction containing a commercial reverse transcriptase (lane 1); molecular weight markers are shown (lanes 3–4). (B) Amplification from cDNA templates generated from the preparations (lanes 3–4) and from a commercial reverse transcriptase control (lanes 1–2); negative controls show no amplification (lanes 5 and 7); DNA ladder shown in lane 6.

### 3.2 Biochmical Signatures and Ancestral Genome Architecture

Nucleic acid extraction from purified preparations revealed a strong predominance of RNA over DNA. Total nucleic acids were resistant to DNase treatment but abolished by RNase digestion, demonstrating that RNA constitutes the primary genetic material of the entities. Robust reverse transcriptase (RT) activity was consistently detected in aliquots containing intact purified entities, whereas matched preparations filtered through 0.2 μm membranes lacked detectable RT activity. The absence of detectable RT activity in filtered supernatants, together with its retention in unfiltered preparations, indicates that the enzymatic activity is associated with cell-sized particles and argues against freely diffusible retroviral components, (Fig. 6).

**Figure 6.**
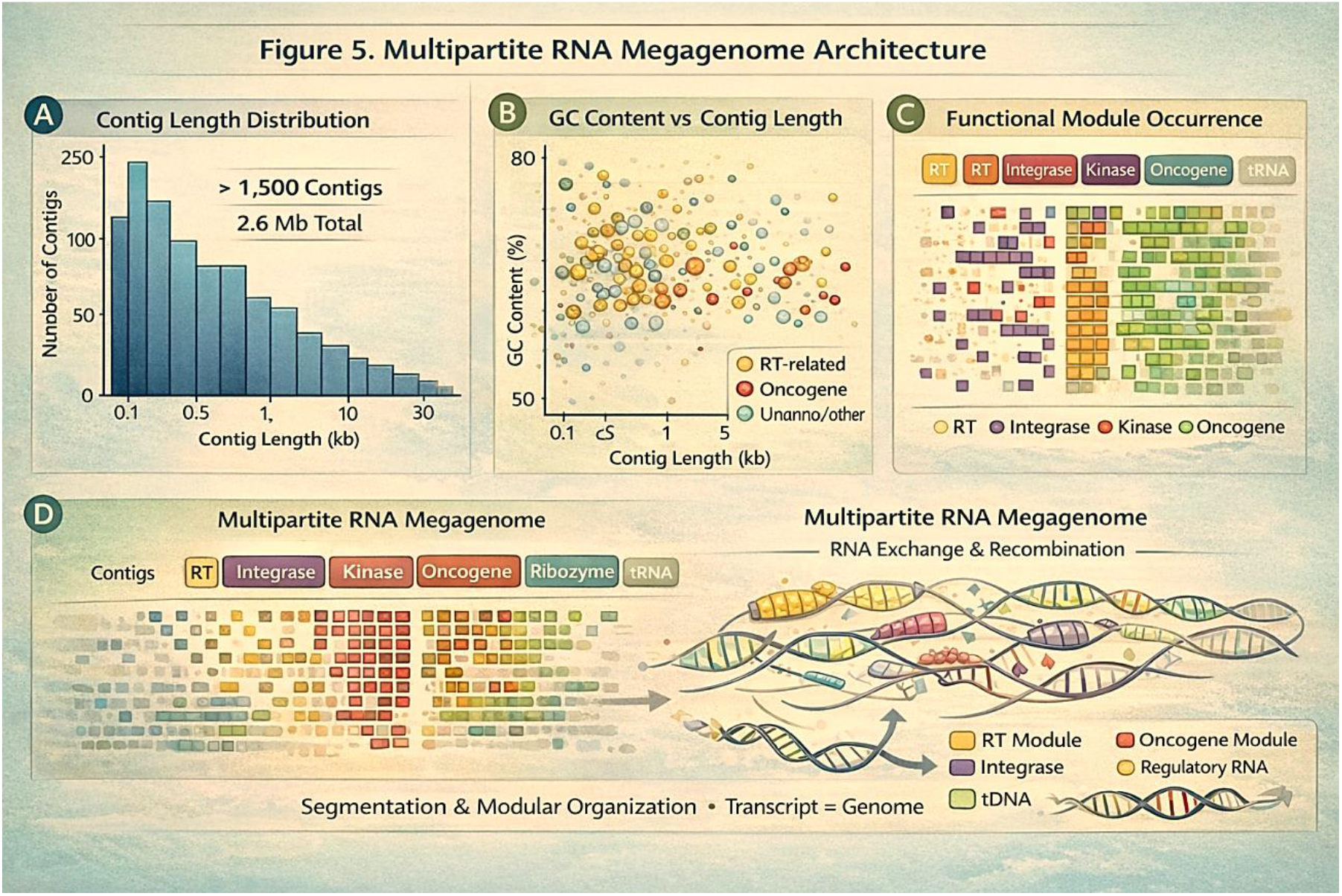
Analysis of RNA-derived sequences from purified preparations of the unicellular agents. (A) Length distribution of RNA-derived contigs reveals a broad, structured size range inconsistent with RNA degradation or linear genome organization. (B) GC content varies widely across contigs, excluding derivation from a single bacterial genome or cellular transcriptome. (C) Detection of functional domains, including reverse transcriptase–related sequences, distributed across multiple non-contiguous contigs. (D) Schematic representation of a modular, non-linear RNA organization observed in the living entities.

### 3.3 RNA analysis indicates a non-linear genetic system associated with Precambrian-like cellular organization

Sequencing of RNA-derived material yielded highly fragmented assemblies that could not be reconciled with conventional linear genome models. Independent sequencing replicates consistently failed to produce a contiguous genome, instead generating non-overlapping contigs distributed across a broad size range. Across datasets, 1,597 contigs were identified, totalling approximately 2.63 Mb, with lengths ranging from ∼100 bp to >30 kb and no dominant size class. GC content varied substantially among contigs, inconsistent with a single clonal bacterial genome or a conventional cellular transcriptome, and indicative of a heterogeneous genetic composition.

Multiple independent observations argue against contamination from known biological sources. The dataset lacked enrichment for bacterial ribosomal RNAs (16S, 23S, 5S), canonical operon structures, and conserved housekeeping genes. Similarly, features characteristic of eukaryotic transcriptomes— including splice-junction enrichment, coordinated tissue-specific expression, and repetitive RNA signatures such as Alu or 7SL-derived elements—were not detected. Sequences encoding reverse transcriptase–related domains were identified, but these were distributed across multiple contigs and were not organized into canonical retroviral genomic architectures (e.g., gag–pol–env), excluding known retroviral systems.

In addition, RNA elements encoding conserved regulatory protein domains, including kinase- and transcription-associated motifs, were reproducibly detected across independent contigs. These sequences occurred within a broader context enriched in mobile element–associated features, including reverse transcriptase–like domains, but were not arranged in recognizable cellular or viral genome structures. Rather than representing degradation or technical artefact, the observed architecture is consistent with a fragmented, non-linear genetic system in which coding elements are distributed across multiple independent RNA units.

While such organization differs from that of known cellular and viral genomes, it is reproducible across preparations and therefore unlikely to reflect stochastic fragmentation. Taken together, the data indicate the presence of an atypical RNA-associated genetic organization linked to the purified entities. The nature and organization of this system remain to be fully resolved (Fig. 6).

The biological and physical characteristics of the entities described here distinguish them from both conventional viruses and known cellular systems. In particular, their size range (1–3 μm) exceeds that of typical filterable viruses, including retroviruses (∼100 nm), and their consistent retention during size-permissive fractionation, together with the absence of reverse transcriptase activity in filtered supernatants, argues against their interpretation as viral particles.

At the same time, these entities are substantially smaller than typical mammalian somatic cells (generally 10–50 μm) and lack the structural organization characteristic of known eukaryotic cell types. Their reproducible isolation as discrete, cell-sized units, coupled with intrinsic enzymatic activity and RNA-dominant genetic material, supports their interpretation as autonomous, biologically active systems rather than subcellular fragments or degenerative cellular debris.

Furthermore, their properties are distinct from those of giant viruses, which are DNA-based systems typically isolated through amoebal hosts and exhibit genomic and replication strategies not observed here. In contrast, the entities described in this study are characterized by RNA-associated genetic organization, reverse transcriptase activity and morphological features comparable to Precambrian microfossils.

Together, these observations support the interpretation that the entities represent a distinct class of small, cell-like biological systems.

## 4. Implications for Classification

The entities described here are not accommodated within established viral or cellular frameworks. Their size (1–3 μm), ultrastructural organization, particle-associated biochemical activity, and RNA-based genetic organization distinguish them from filterable viruses, including retroviruses, while their dimensions and morphology exclude known cellular systems. Together, their Precambrian-like morphological features and associated biochemical properties support their interpretation as a distinct class of small, autonomous cell-like biological systems and warrant consideration of a dedicated taxonomic framework.

## 5. Discussion

The identification of living unicellular entities displaying morphological features reminiscent of Precambrian microfossils within mammalian tissues raises questions that challenge prevailing assumptions in evolutionary biology and cellular organization. Rather than representing fossilized relics or artifacts of preservation, these entities are observed as metabolically active systems reproducibly recovered from independent specimens.

The Precambrian Eon, extending from the origin of life to approximately 541 million years ago, was dominated by microbial ecosystems characterized by extensive structural and organizational diversity. During this interval, biological systems exhibited a wide range of configurations, including organic-walled vesicles, cleavage-like internal partitioning, and early forms of developmental organization.

The entities described here, typically measuring 1–3 μm in diameter, display comparable features, including organic wall–bounded morphology, internal compartmentalization consistent with cleavage-like organization and staged developmental progression. Their correspondence with acritarchs, Doushantuo embryo-like forms and other Proterozoic microfossils extends beyond superficial resemblance to include patterns of organization historically interpreted as indicative of early cellular systems (Xiao et al., 2004; Butterfield, 2018; Javaux et al., 2010; Loron et al., 2019).

The degree of ultrastructural correspondence indicates that fundamental architectural features characteristic of Precambrian microfossils are present in these entities. Previous work has shown that morphological complexity in early microfossils reflects biological strategies that are not readily mapped onto known lineages (Javaux, 2019; Knoll et al., 2006). In this context, the Precambrian-like morphology reflects a defined biological strategy of cellular organization, revealing in living systems biological architectures previously inferred only from the fossil record.

Initial attempts at genomic characterization yielded results that do not conform to conventional DNA-based cellular or viral systems. Sequencing of RNA-derived material consistently produced fragmented assemblies that could not be reconstructed into a contiguous genome and lacked canonical features of bacterial, eukaryotic, or viral genomic organization. The reproducibility of this pattern across independent preparations indicates that it reflects an intrinsic property of the system rather than technical artifact.

In the context of the observed morphology, this genomic organization is consistent with models in which genetic information is distributed across multiple RNA elements rather than encoded within a single linear genome. The presence of intrinsic reverse transcriptase activity further supports RNA-associated replication mechanisms. Such features are compatible with theoretical models of early biological systems in which RNA fulfilled both informational and catalytic roles (Gilbert, 1986; Joyce, 2002; Higgs and Lehman, 2015; Saito, 2022), although their precise relationship to the entities described here remains to be defined.

The biological relevance of these entities is supported by multiple independent lines of evidence, including reproducible purification, electron microscopy, and particle-associated enzymatic activity. Their consistent isolation from independent specimens and stability under experimental conditions argue against stochastic contamination and support their interpretation as coherent biological systems. Similar criteria—reproducibility, structural integrity, and contextual consistency—have been emphasized in the evaluation of ancient microfossils and their biogenicity (Donoghue, 2020; Schopf, 1993; Brasier et al., 2006).

Collectively, the combination of Precambrian-like morphology, RNA-associated genetic features and intrinsic enzymatic activity indicates the presence of a form of unicellular organization that is not readily accommodated within established biological frameworks. The entities described here are clearly distinct from both viral particles and conventional cellular systems. Their size, structural organization, and intrinsic biochemical activity differentiate them from filterable viruses, while their dimensions and morphology exclude classification as known eukaryotic cells. This coherent biological profile supports their interpretation as autonomous cell-like systems rather than variants of established categories. Comparable challenges in classification arise where morphological disparity exceeds current taxonomic frameworks (Knoll et al., 2006; Butterfield, 2018).

Taken together, these observations support a connection between palaeobiological structures and contemporary biological systems. If cellular architectures of Precambrian microfossils are recapitulated in such entities, then aspects of early biological organization may remain accessible to direct investigation. This perspective invites a reassessment of how ancient cellular designs are interpreted and suggests that foundational principles of early evolution may persist in forms that have not yet been fully recognized.

In this context, the purpose of the present manuscript is to invite expert assessment focused specifically on the morphological characteristics of these entities within a palaeobiological framework. While limited biochemical observations are included, these serve to demonstrate that the agents described are not inert relics, but biologically active systems. A more comprehensive analysis of their biochemical and molecular properties will be addressed in subsequent dedicated studies.

More broadly, these findings suggest a potential expansion in the scope of Precambrian research. Traditionally centred on fossil evidence and hypothetical reconstruction, the field may extend toward the investigation of biological mechanisms that persist in living systems. In this perspective, the Precambrian is not only a record of early life, but may also contribute to understanding fundamental processes relevant to modern biology. Such a view is consistent with increasing recognition that deeply conserved cellular mechanisms and genomic architectures continue to inform present-day biological function and disease processes (Lane, 2015; Knoll and Nowak, 2017).

## Conclusion

The findings presented here document, for the first time, the existence of living cell-like entities displaying ultrastructural features closely comparable to Precambrian microfossils, in association with RNA-dominant genetic material and intrinsic reverse transcriptase activity. The reproducible identification of these entities in present-day mammalian systems, together with their defined morphology and biochemical properties, establishes them as biologically active agents rather than inert structures.

The combination of Precambrian-like cellular architecture and distinctive RNA biochemistry distinguishes these entities from known cellular and viral forms and indicates the presence of a type of biological organization not currently accommodated within established classification frameworks. On this basis, consideration of an expanded or revised taxonomic framework is warranted.

These observations underscore the foundational role of Precambrian palaeobiology. Morphological frameworks developed through decades of research on the fossil record—particularly on acritarchs, Doushantuo embryo-like forms and related microfossils—provide the essential basis for recognizing and interpreting these entities. In this sense, the present findings directly build upon the contributions of palaeontological research.

By demonstrating that cellular entities previously known only from the Precambrian fossil record can be identified in living systems, this work establishes a direct link between palaeobiological observations and contemporary biology, providing a new framework for investigating early cellular organization in modern biological contexts. Biological strategies first inferred from ancient microfossils are now directly observable in active biological systems.

## Author Contributions (CRediT Statement)

**Lusi Elena Angela**: Conceptualization; Methodology; Investigation; Formal analysis; Data curation; Writing – original draft; Writing – review & editing; Supervision; Project administration.

**Federico Caicci**: Methodology; Investigation; Validation.

**Quartuccio Marco**: Resources; Data curation.

**Claudia Rifici:** Investigation; Data curation; Validation; Writing – review & editing.

## Notes

https://doi.org/10.6084/m9.figshare.31802113

### Competing Interest Statement

The authors have declared no competing interest.

